# Neutralizing antibodies elicited in sequentially immunized macaques recognize V3 residues on altered conformations of HIV-1 Env trimer

**DOI:** 10.1101/2024.07.31.605918

**Authors:** Andrew T. DeLaitsch, Jennifer R. Keeffe, Harry B. Gristick, Juliet A. Lee, Wenge Ding, Weimin Liu, Ashwin N. Skelly, George M. Shaw, Beatrice H. Hahn, Pamela J. Björkman

## Abstract

Eliciting broadly neutralizing antibodies that protect against diverse HIV-1 strains is a primary goal of AIDS vaccine research. We characterized Ab1456 and Ab1271, two heterologously-neutralizing antibodies elicited in non-human primates by priming with an engineered V3-targeting SOSIP Env immunogen and boosting with increasingly native-like SOSIP Envs derived from different strain backgrounds. Structures of Env trimers in complex with these antibodies revealed V3 targeting, but on conformational states of Env distinct from the typical closed, prefusion trimeric SOSIP structure. Env trimers bound by Ab1456 adopted conformations resembling CD4-bound open Env states in the absence of soluble CD4, whereas trimers bound by Ab1271 exhibited a trimer apex-altered conformation to accommodate antibody binding. The finding that elicited antibodies cross-neutralized by targeting altered, non-closed, prefusion Env trimer conformations provides important information about Env dynamics that is relevant for HIV-1 vaccine design aimed at raising antibodies to desired epitopes on closed pre-fusion Env trimers.

**Highlights:**

- Sequential immunization regimen elicits V3 antibodies targeting non-closed Envs
- Cryo-EM structures reveal recognition of multiple Env conformational states
- Neutralization by elicited antibody does not require antibody-virus preincubation

**Graphical Abstract:** 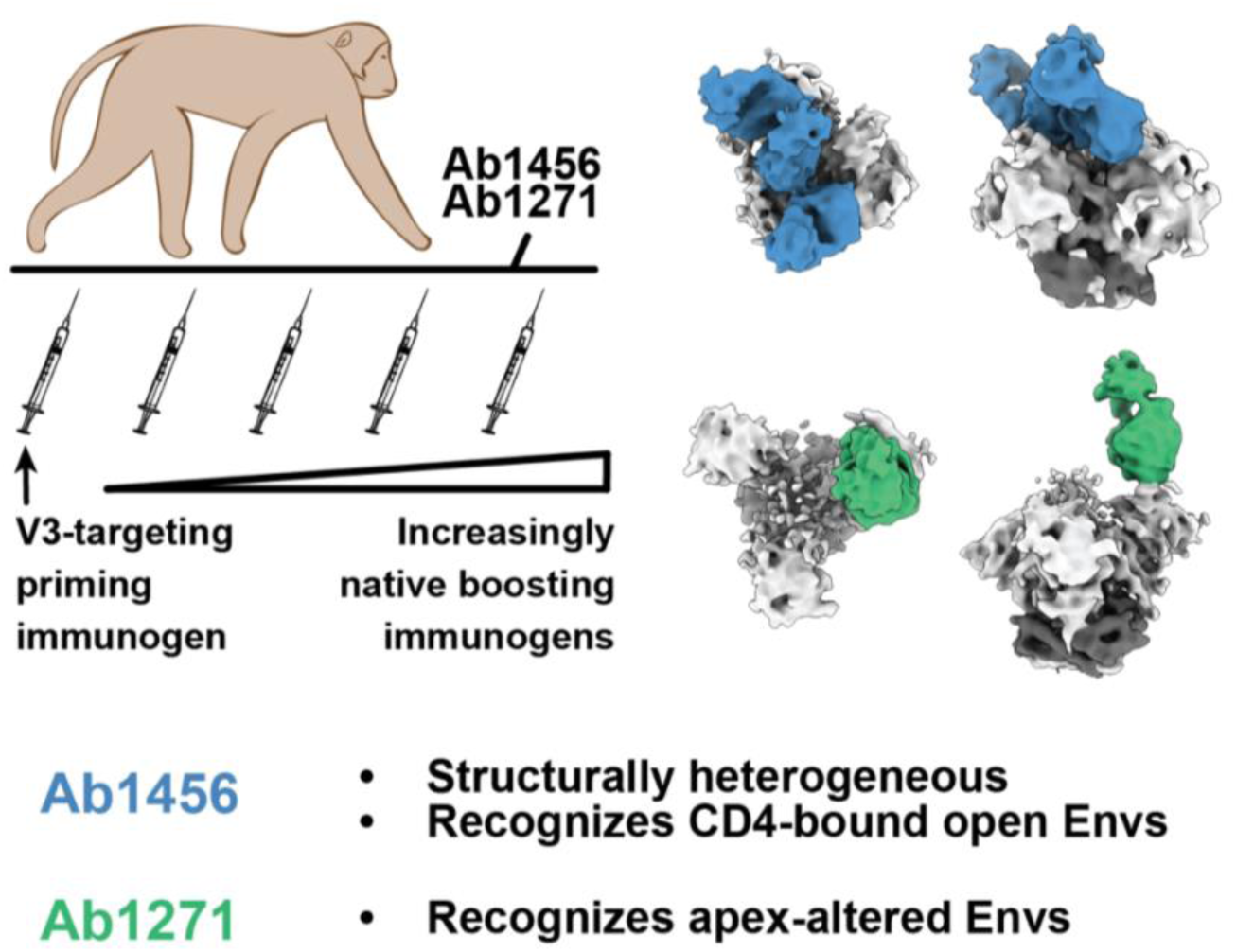

## Introduction

The HIV-1 envelope glycoprotein (Env) on the virion surface is responsible for fusing viral and host cell membranes during infection.^1^ Env, a heavily glycosylated trimer of gp120-gp41 heterodimers, functions via a dynamic mechanism initiated upon engaging one or more copies of the host cell receptor CD4.^1^ CD4 binding leads to open conformational states of Env trimer in which gp120s undergo an outward rotation,^2^ and protomers bound by CD4 exhibit a large-scale rearrangement in the V1V2 region of gp120 that exposes the binding site for the HIV-1 co-receptor, CCR5 or CXCR4.^3^ This conformation can be described as a CD4-bound open state, as it is typically observed in the presence of CD4 or a CD4 mimetic small molecule.^3–6^ After co-receptor binding, gp41-mediated fusion of viral and host cell membranes allows the HIV-1 genetic material to enter the host cell to establish an infection.^1^

Neutralizing antibodies against HIV-1 solely target Env, where they act to prevent fusion of the viral and host cell membranes. As such, Env comprises a key target of HIV-1 vaccine strategies.^7,8^ Because the rapid mutation rate of HIV-1 creates high levels of sequence diversity both within and between hosts, an effective prophylactic vaccine will need to induce broadly neutralizing antibodies (bNAbs) capable of recognizing not one, but many, circulating strains.^7,8^ bNAbs isolated from people living with HIV-1 target conserved features on Env, including the CD4 binding-site (CD4bs) and the V3-loop at the Env trimer apex involved in co-receptor binding.^9^ Structures of bNAb-Env complexes mainly reveal targeting of pre-fusion closed Env trimers^3^ with the exception of b12, one of the first characterized bNAbs.^10^ b12 targets an Env conformation in which the gp120s undergo an outward rotation, but V1V2 remains on top of V3, thereby occluding access to the co-receptor binding site and distinguishing this occluded-open conformation from the CD4-bound open conformation.^11–13^

In an attempt to produce bNAbs in wildtype animals with polyclonal antibody repertoires, we previously described an immunization protocol that involved priming with a V3 germline-targeting Env immunogen^14^ followed by sequential boosting with increasingly “native” Env trimers.^15^ After boosts 3 and 4, we isolated heterologous, but weakly neutralizing, monoclonal antibodies (mAbs) from immunized non-human primates (NHPs). Seven of nine NHP mAbs elicited after the prime-boost regimen targeted the V3-glycan patch, as demonstrated by competition with the V3-targeting bNAb 10-1074.^15^ However, the remaining two mAbs, Ab1303 and Ab1573, competed with the CD4bs bNAb 3BNC117^15^ and were shown by single-particle cryo-electron microscopy (cryo-EM) to target the CD4bs of Env trimers in an occluded-open, rather than a CD4-bound open, conformation.^12^

Here, we report cryo-EM structures of two of the 10-1074-competing NHP mAbs from the prime-boost sequential immunization regimen^15^ in complex with stabilized, soluble Env trimeric ectodomains (SOSIPs).^16^ In common with Ab1303 and Ab1573, which were also elicited in this immunization regimen, neither of the V3-targeting NHP mAbs recognized the pre-fusion closed Env conformation. Instead, Ab1456 interacted with the V3-epitope on Env trimers adopting various CD4-bound open conformations although neither soluble CD4 (sCD4) nor a CD4 mimetic was included in the complex. The other antibody, Ab1271, also interacted with the V3 region of an Env trimer but recognized a distinct Env conformation in which V1V2 was displaced but the gp120s did not exhibit outward rotation. The discovery of apparently preferential targeting of non-closed Env conformations by antibodies elicited by sequential immunization with SOSIP-based immunogens has important implications for HIV/AIDS vaccine design.

## Results

### Ab1456 and Ab1271 are heterologously-neutralizing mAbs elicited in sequentially immunized NHPs

Ab1456 and Ab1271 were isolated from NHPs after sequential immunizations with engineered or wildtype SOSIP-based immunogens designed to target the V3-glycan patch on the gp120 subunit of Env and characterized as weak, but heterologously-neutralizing mAbs.^14,15^ NHPs were primed with RC1-4fill, a low affinity V3-glycan patch germline-targeting immunogen conjugated to virus-like particles (VLPs) using the SpyCatcher-SpyTag system.^17,18^ RC1-4fill is a modification of the clade A BG505-based 11MUTB SOSIP immunogen,^19^ in which the N156_gp120_ glycan was removed (N156Q) and potential N-linked glycosylation sites (PNGSs) to block BG505 strain-specific responses to an immunodominant glycan hole in the vicinity of residue 241_gp120_^20^ were added. A series of boosts consisting of VLPs presenting 11MUTB-4fill,^15^ a clade B B41^21^ or B41-5MUT,^15^ a mosaic of a clade B (AMC011)^22^ and clade C (Du422)^23^, and a mosaic of consensus Envs from Group M and clade C (ConM/ConC)^24,25^ were given to try to shepherd antibody responses towards broader reactivities and avoid strain-specific responses (Figure 1A).

**Figure 1.**
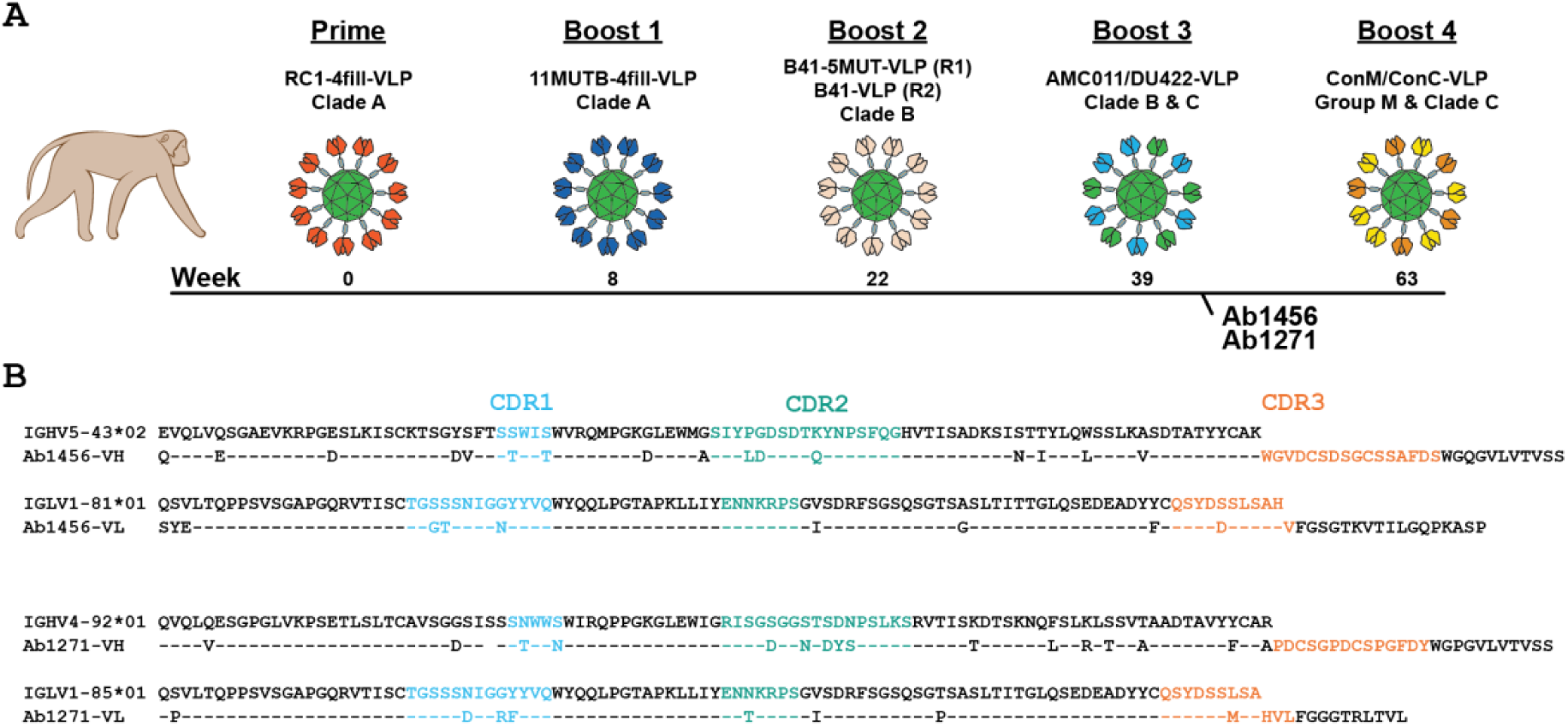
Characterization of Ab1456 and Ab1271. (A) Schematic describing the sequential immunization of NHPs that gave rise to Ab1456 (Regimen 1 in Boost 2; R1) and Ab1271 (Regimen 2 in Boost 2; R2).^15^ Ab1456 was isolated from NHP DGJI, and Ab1271 was isolated from NHP T15. (B) Alignments of Ab1456 and Ab1271 to their presumptive germline VH gene precursors, as identified by IMGT/V-QUEST.^27,28^ CDRs are defined according to Kabat.^29^

Ab1456 and Ab1271 were isolated after the third boost in the same two NHPs as the Ab1573 and Ab1303 CD4bs mAbs (Figure 1A).^15^ Ab1456 was derived from macaque IGHV5-43*02 and IGLV1-81*01 germline V gene segments, exhibiting 14.3% (heavy chain; HC) and 20.2% (light chain; LC) amino acid changes due to somatic hypermutation. Ab1271, derived from the IGHV4-92*01 and IGLV1-85*01 germline V gene segments, exhibited 15.5% (HC) and 7.4% (LC) changes from somatic hypermutation (Figure 1B). Of note, there is a one-residue deletion in the HC framework region 1 (FWRH1) of Ab1271. In contrast to the Ab1573 and Ab1303 CD4bs mAbs, Ab1456 and Ab1271 each competed with the V3 bNAb 10-1074, suggesting on-target binding specificities for these mAbs.^15^ Both mAbs displayed heterologous neutralization when tested against a panel of 19 pseudoviruses including the 12-strain global HIV-1 panel^26^ and two SHIVs, neutralizing 6 of 19 (Ab1456) or 14 of 19 (Ab1271) HIV-1 pseudoviruses with IC_50_ values <100 µg/mL.^15^ While neutralization potencies were generally weak, Ab1271 in particular exhibited breadth, neutralizing all viruses tested at IC_50_ values <500 µg/mL.

### Ab1456 recognizes CD4-bound open Env trimer conformations

For structural studies of Ab1456 recognition of Env, we formed Fab-Env complexes using a chimeric SOSIP Env containing a gp120 derived from JRCSF.JB, an HIV-1 strain that was potently neutralized by Ab1456,^15^ and a gp41 derived from BG505.^30^ Fab-SOSIP complexes isolated by size-exclusion chromatography (SEC) were used for EM analysis. Initial processing in cryoSPARC^31^ yielded a 6.6 Å resolution structure that showed targeting of the trimer apex of an open Env conformation with an apparent stoichiometry of one Fab per trimer (Figure 2A,B). In this structure, two of three protomers in the trimer appeared to adopt a CD4-bound open conformation as indicated by rearranged V1V2 densities, while the third protomer exhibited an outward rotation, but lacked the V1V2 rearrangement.^32^ Overall, this state of the Env trimer resembled the conformation of HT2, a SOSIP heterotrimer in complex with two, rather than three, copies of soluble CD4;^6^ hence, we refer to this Ab1456-bound Env conformation as HT2-like, noting, however, that the Ab1456-Env structure was determined in the absence of CD4 and with a homotrimeric SOSIP (Figure 2A,B). Unexplained density was present at the trimer apex (Figure S1A), suggesting that this consensus structure included particles from distinct 3D classes. Indeed, 3D classification in RELION^33^ revealed extensive heterogeneity. Structural classes were identified that differed in the conformational state of the trimer, the number of bound Fabs per trimer, the relative positioning of the bound protomers, and the approach angles of the Fabs (Figure 2). Sorting of approximately 80,000 particles allowed us to determine eight structural classes with resolutions ranging from 8.8 Å to 14 Å (Figure S2). Given the high degree of heterogeneity, additional states could exist. In addition, imperfect separation of particles may bias some of the reported structures.

**Figure 2.**
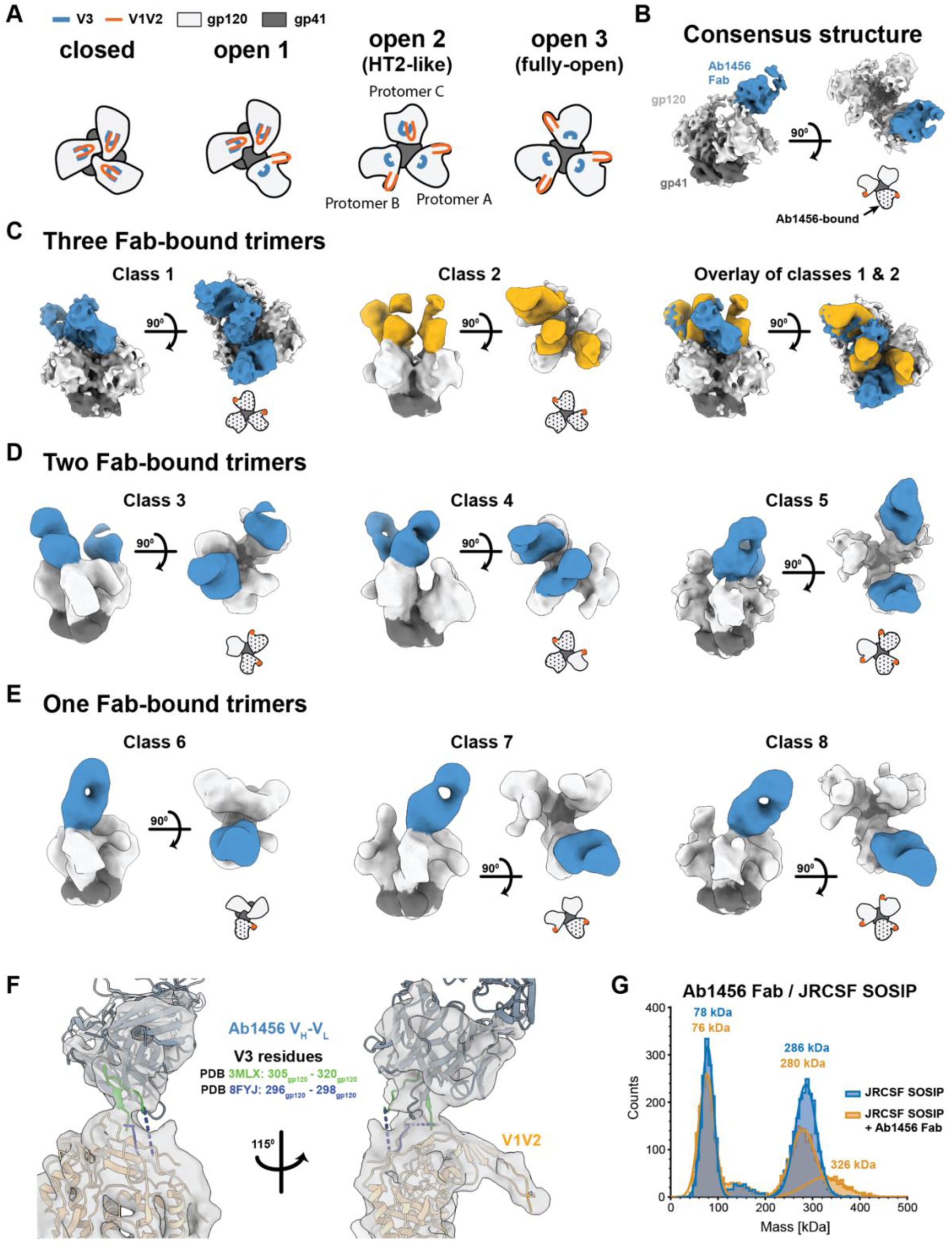
Ab1456 Fab binds open forms of HIV-1 Env. (A) Schematic of top-down views of Env trimers showing a potential pathway from a closed (left) to a fully-open (right) trimer. The three open Env states were identified in structural classes with bound Ab1456 Fab(s). (B) Results of a non-uniform refinement of Ab1456 Fab-bound JRCSF SOSIP particles prior to 3D classification. A schematic representation of the top-down view of the trimer conformation is shown in this and subsequent panels with the Ab1456-bound protomer(s) indicated by a black dotted pattern and displaced V1V2 loop(s) indicated by orange dot(s). (C) Left and middle: Two structural classes showing three Ab1456 Fabs bound. Right: An overlay of the two structural classes. (D) Three structural classes showing two Ab1456 Fab-bound trimers. (E) Three structural classes showing one Ab1456 Fab-bound trimers. (F) Analysis of the Ab1456 epitope. Protomer A from PDB 8FYJ (HT2 Env heterotrimer complexed with two CD4s) and 3MLX (human mAb 3074 complexed with a V3 peptide; ordered residues 305-320) were independently docked into the density corresponding to protomer A and Ab1456 Fab / V3 peptide in the consensus map, respectively. (G) Mass photometry of JRCSF SOSIP (blue) and JRCSF SOSIP in complex with Ab1456 Fab (gold).

Two different three Fab-bound HT2-like trimer classes were identified. In these classes, the Fab bound to protomer B was wedged either in front, or behind, of the Fab bound to protomer C, resulting in distinct angles of approach (Classes 1 and 2) (Figure 2C).

Multiple structural classes of Env trimers with two bound Fabs were also found. Two distinct classes of two Fab-bound HT2-like trimers were identified with the Ab1456 Fabs binding to different configurations of the Env A, B, and C protomers. In one configuration, both Fab-bound protomers (protomers A and B) adopted the CD4-bound open conformation (Class 3) (Figure 2D).

In another structural class, both the protomer that lacked apparent V1V2 rearrangements (protomer C) and an adjacent protomer in the CD4-bound open conformation (protomer A) exhibited bound Ab1456 Fabs (Class 4) (Figure 2D). Another structural class showed two Ab1456 Fabs bound to a fully-open trimer in which all three protomers displayed V1V2 rearrangements (Class 5) (Figure 2D).

In Env structures exhibiting a single bound Fab, three different trimer conformational states were identified (Figure 2A,E). In one state, only the Fab-bound protomer adopted a CD4-bound open conformation (as defined by a V1V2 rearrangement), and the other protomers exhibited neither an outward gp120 rotation nor V1V2 rearrangement (Class 6) (Figure 2E). Another class was found in which the Env trimer adopted an HT2-like state, and only protomer A was bound (Class 7) (Figure 2E). Finally, a structure of a single Ab1456 Fab bound to a trimer in which all three protomers adopted a CD4-bound open conformation was identified (Class 8) (Figure 2E).

At the resolutions of our EM structures, we are mostly limited to analyzing the Ab1456 epitope through docking of previously-determined EM and X-ray structures. Docking of protomer A from the HT2 trimer (Env heterotrimer bound by 2 copies of soluble CD4; PDB 8FYJ) into the consensus structure revealed qualitative agreement with protomer A density in the EM map (Figure S1B). The docked protomer showed apparent Ab1456 targeting of V3 residues that were not built in the HT2 structure as a consequence of being disordered in PDB 8FYJ. To account for additional Ab1456 and Env V3 density in the Ab1456-JRCSF Env structure, we docked a crystal structure of the human mAb 3074 in complex with a V3 peptide spanning gp120 residues 301-324 (PDB 3MLX; residues 305-320 were ordered in the crystal structure). We chose this peptide/V3-antibody structure because 3074 neutralizes viruses in common with Ab1456 (e.g., both neutralized JRCSF, 6535.3, and X1632 to a greater extent than other strains^15,34^) and preferentially binds Env in the presence of CD4.^35^ While these properties may indicate that 3074 and Ab1456 make similar contacts with Env, such interactions could be mediated by different antibody features (e.g., different complementarity-determining regions, different specific Fab-Env interactions, different Fab binding orientations, etc.). The docked Fab-bound V3 peptide fit the EM density well, providing support for the interpretation that Ab1456 and mAb 3074 contact similar V3 residues that are exposed on open states of the trimer (Figure 2F). Ab1456 targeting of the V3 epitope is consistent with it competing for Env binding with the human V3-directed bNAb 10-1074.^15^

To further characterize the stoichiometry of Ab1456 Fab binding to Env trimer, we performed mass photometry, a technique that detects binding interactions in solution via mass measurements of individual molecules.^36,37^ Unlike the Ab1456-JRCSF SOSIP complex used for cryo-EM, the Fab-Env sample for mass photometry was not purified by SEC, and mass photometry was performed at a more dilute final concentration (diluted from ∼1 mg/mL to <1 µg/mL for measurement) than what was imaged by cryo-EM (∼1.1 mg/mL). By mass photometry, the Ab1456-JRCSF sample showed particles with increased mass relative to the trimer alone control (Figure 2G). Although distinct populations could not be unambiguously identified, the masses of the complexes were consistent with a mixture of Fab-Env particles containing either 0, 1, 2, or 3 bound Fabs per trimer (Figure 2G).

### Ab1271 recognizes a conformation distinct from pre-fusion closed Env trimers

We next focused on Ab1271, which exhibits a broader neutralization profile than Ab1456.^15^ For these structural studies, we formed Fab-Env complexes using a chimeric SOSIP with a gp120 derived from 6535.3, a tier 1 virus that was potently neutralized by Ab1271,^15^ and a gp41 derived from BG505.^30^ Cryo-EM analysis revealed trimers that were not complexed with Ab1271 Fab, yielding a 4.6 Å structure of the unliganded 6535.3 SOSIP trimer (Figure 3A, Figure S3). Although the 6535.3 Env conformation generally resembled a typical closed prefusion trimer in that gp120s were not outwardly rotated and the V1V2 regions were not displaced to the trimer sides as seen in the CD4-bound open Env conformation,^11,32,38^ the trimer apex in 6535.3 Env differed from those of BG505 and other SOSIP Env structures. Docking of a closed BG505 SOSIP (PDB 6UDJ) into the 6535.3 SOSIP density showed differences in the presumptive locations of the V1V2 regions at the trimer apex for two of the three 6535.3 protomers, and density for V1V2 on the third protomer was not observed (Figure 3A).

**Figure 3.**
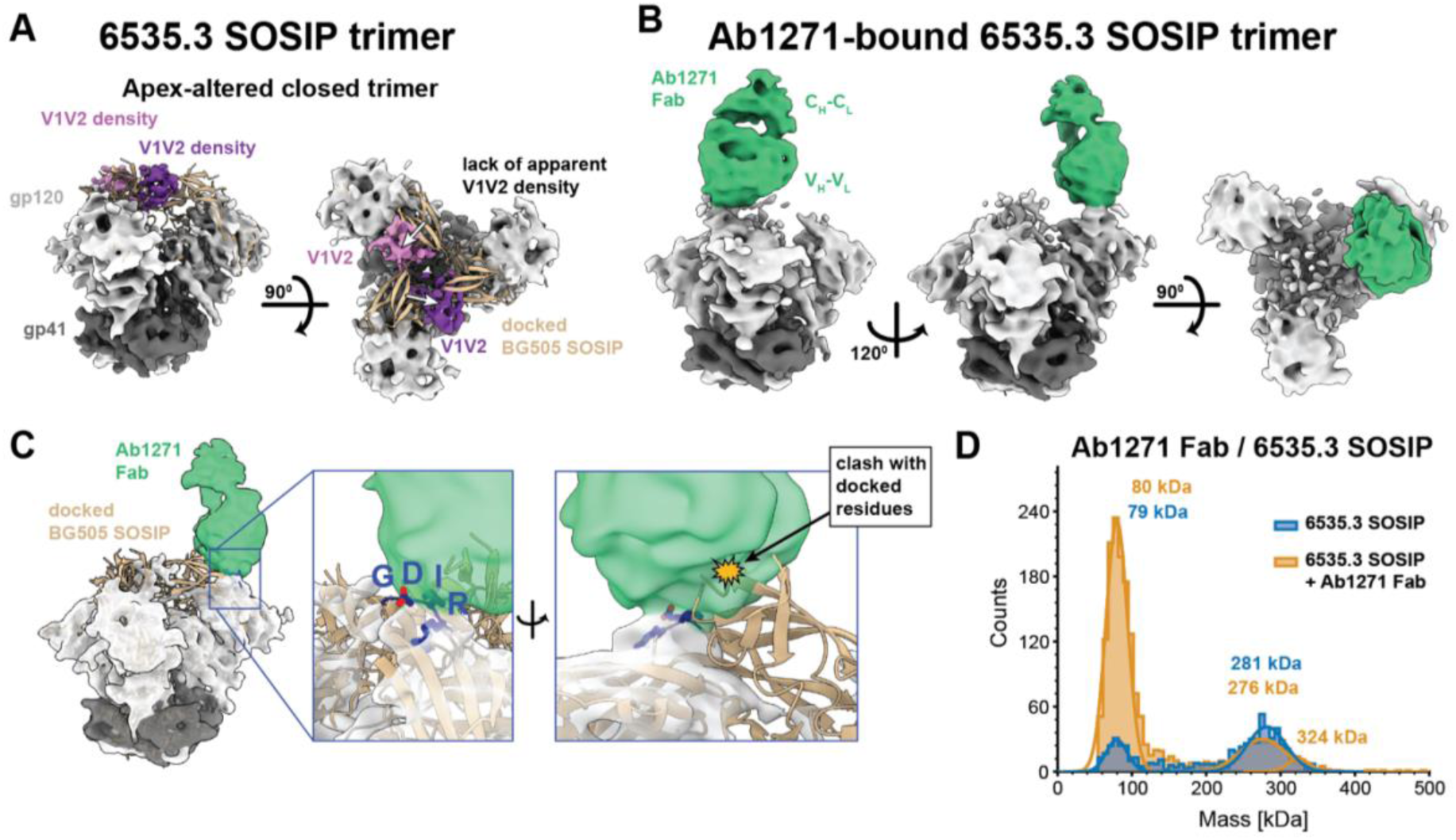
Ab1271 recognizes a closed 6535.3 Env trimer with an altered apex. (A) Side and top-down views of a structural class of unliganded 6535.3 SOSIP. BG505 SOSIP (PDB 6UDJ) (tan cartoon representation) was docked into 6535.3 density. Presumptive V1V2 density for two of the three 6535.3 protomers (pink and purple) is shifted relative to the BG505 V1V2, as indicated by white arrows in the top-down view on the right. (B) A structural class of Ab1271-Env complex with a single copy of Ab1271 Fab bound to the 6535.3 SOSIP. (C) Left: Ab1271-6535.3 complex. Middle: Close-up of Fab-Env interaction showing apparent targeting of Ab1271 Fab towards the V3 GDIR motif (G324_gp120_ – R327_gp120_). Right: Close-up of Fab-Env interaction showing apparent clash with V1V2 residues of the docked BG505 SOSIP. (D) Mass photometry of 6535.3 SOSIP (blue) and 6535.3 SOSIP in complex with Ab1271 Fab (gold).

In addition to the unbound 6535.3 trimer structure, we determined a 6.3 Å structure of a 6535.3 SOSIP bound by a single Ab1271 Fab (Figure 3B). In this structure, the Env portion of the density closely matched the density for the unbound 6535.3 SOSIP (Figure 3A,B), with Ab1271 interacting with the V3 region of the protomer lacking apparent V1V2 density. The apex of a docked closed BG505 trimer (PDB 6UDJ) clashed with density corresponding to the Ab1271 Fab, suggesting that V1V2 must be in a position distinct from its position in BG505 in order to accommodate Ab1271 binding (Figure 3C). Unlike the occluded-open Env conformations found in the Ab1303-Env and Ab1573-Env complex structures^12^ or the CD4-bound open structures with Ab1456 (Figure 2), the Env trimer in the Ab1271-Env complex did not exhibit an outward rotation of its gp120s, in common with conventional pre-fusion closed SOSIP trimer structures.^3^ However, as the form of the Env trimer in the Ab1271-6535.3 SOSIP complex is distinct from previously-determined closed trimer structures,^3^ we refer to its conformation as apex-altered closed. While low resolution limited our analysis of antibody epitope details, the docked structure revealed apparent Ab1271 targeting at or near the conserved GDIR motif within the V3 loop of gp120^39^ (gp120_324-327_) (Figure 3C).

To further investigate the stoichiometry of Ab1271 Fab binding to the Env trimer, we evaluated Ab1271 Fab-6535.3 SOSIP complex formation using mass photometry. Similar to the JRCSF SOSIP, the 6535.3 SOSIP also included multiple populations, predominantly corresponding to SOSIP trimers and protomers (Figure 3D). Although a majority of the trimers remained unbound in the presence of Ab1271 Fab, a small shoulder at ∼324 kDa was present in the experimental histogram, consistent with a population of one Fab-bound trimers (Figure 3D). As mass photometry experiments were conducted at more dilute final concentrations (diluted from ∼1 mg/mL to <1 µg/mL) compared to cryo-EM (∼1.9 mg/mL), this result suggests a weak affinity and/or fast off-rate of Ab1271 Fab for the 6535.3 SOSIP trimer.

### Antibody-virus pre-incubation is not necessary for in vitro neutralization by Ab1456

Based on the cryo-EM structures of the weakly neutralizing NHP mAb Ab1456 that revealed recognition of open Env conformations, we reasoned that Ab1456 targeting could be limited by the conformational availability of the epitope on virion-bound Env trimers. Standard TZM-bl neutralization assays provide a time window, typically 1 hour, in which an antibody is incubated at 37 °C with virus prior to the addition of target cells.^40,41^ We hypothesized that this incubation could allow sampling of open trimer conformations, which could then be captured, permitting antibody binding to virion Envs to achieve neutralization. The antibody-virus co-incubation step is distinct from how antibodies neutralize HIV-1 in vivo, where antibodies and viruses are not pre-incubated in a small volume and where there might only be a limited time window for an antibody to recognize Env on a virus prior to encountering a target cell. We therefore reasoned that pre-incubation of antibody and virus might artificially inflate the neutralization potencies of antibodies that target an epitope on an open Env trimer.

To test this possibility, we compared the 50% inhibitory concentrations (IC_50_s) of Ab1456 and other weakly and broadly neutralizing mAbs in the standard TZMbl assay, in which virus and antibodies were preincubated for 1 hour,^41,42^ and in a modified assay, in which antibodies were first added to the cells followed by virus addition in a separate step. Selecting a set of both sensitive and more resistant viruses,^15^ we found that the neutralization potencies of Ab1456 were very similar regardless of whether the standard or modified assay was used (Figure 4A). This was true for HEK293T-derived pseudoviruses and replication-competent simian-human immunodeficiency viruses (SHIVs) as well as a SHIV challenge stock that was propagated in rhesus macaque peripheral blood mononuclear cells (PBMCs).^15,43^ Preincubation also had no effect on the neutralization potencies of other antibodies known to target open Env conformations, such as the CD4-induced antibody 17b^2^ and the linear V3 mAb 3074^44^ (Figure 4B). The slopes of the 17b and 3074 neutralization curves, like those of Ab1456, were more shallow than the slopes of potent bNAbs, such as the V3 bNAb 10-1074^45^ and the CD4bs bNAb VRC01^46^ that recognize closed Env conformations (Figure S4). Since shallow dose-response curves are associated with less favorable therapeutic potentials,^47^ these results imply limited in vivo protection efficacy for Ab1456.

**Figure 4.**
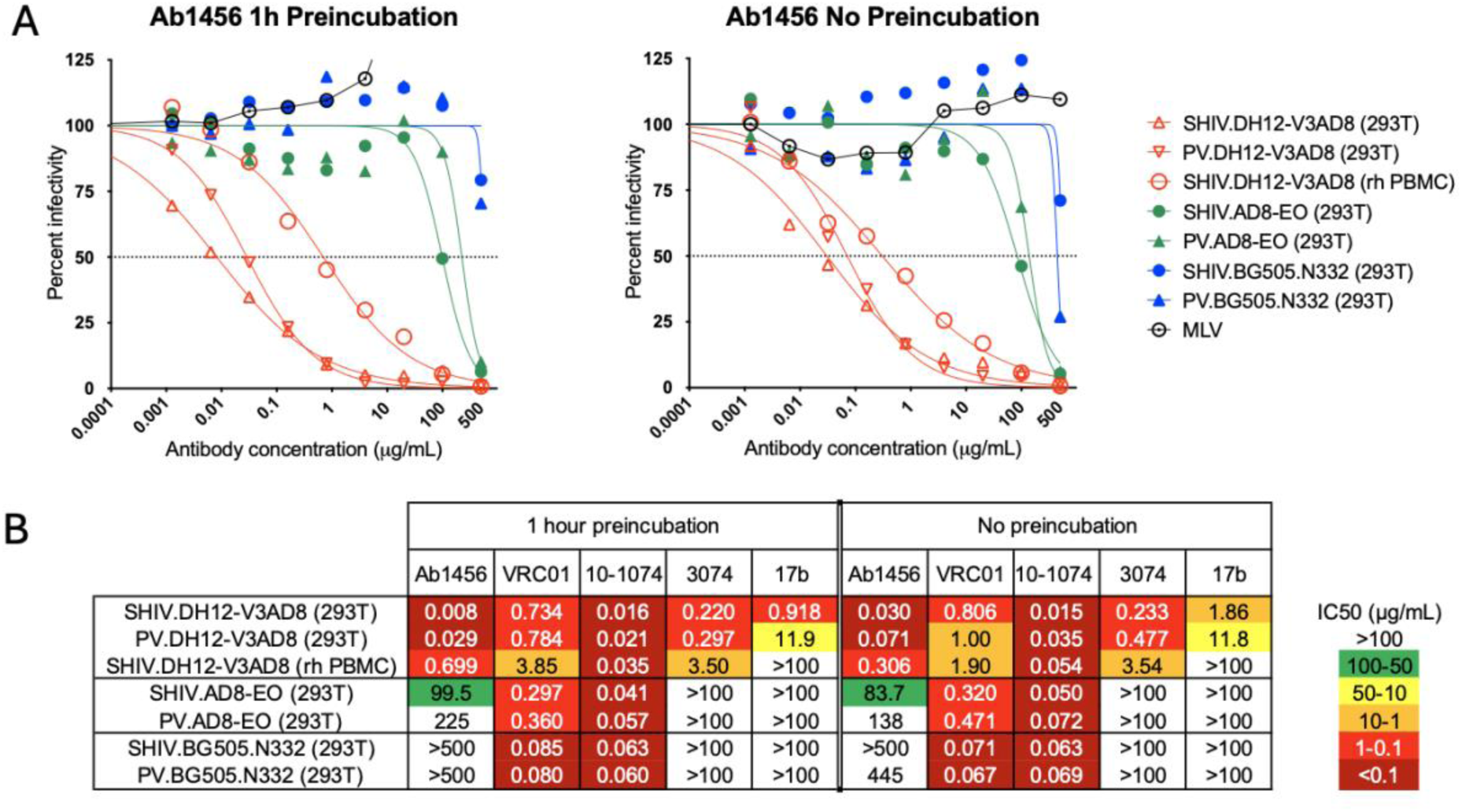
Preincubation of antibody and virus during in vitro neutralization does not affect neutralization potency. (A) Sensitivity of viruses expressing the DH12-V3AD8 (SHIV_DH12-V3AD8_; red), AD8-EO (SHIV_AD8EO_; green) and BG505.N332 (blue) Envs to neutralization by Ab1456 using a TZMbl assay with (left) and without (right) a 1 hour antibody and virus preincubation step. Neutralization curves are shown for pseudovirus (PV) and replication-competent forms of SHIVs derived either by HEK293T transfection (293T) or following propagation in rhesus macaque PBMCs (rh PBMC). Dotted lines indicate 50% reduction in virus infectivity. (B) Sensitivity of the viruses shown in A (listed on the left) to neutralization by other mAbs (listed on top) with (left panel) and without (right panel) a 1 hour antibody and virus preincubation step. 50% inhibitory concentrations (IC_50_) are shown in μg/mL (coloring indicates relative neutralization potency).

## Discussion

The results reported here provide potentially valuable information for the design of immunogens to elicit bNAbs that target closed, prefusion HIV-1 Env trimers rather than antibodies such as Ab1456, which recognize non-closed Env conformations, neutralize heterologous HIV-1 strains only weakly, and lack breadth against difficult-to-neutralize tier 2 HIV-1 strains. Here, we show from cryo-EM structures that our RC1-based prime-boost regimen successfully elicited antibodies against the V3 glycan patch, as previously predicted from competition with the human V3 bNAb 10-1074^15^. As there are no available structures showing 10-1074 recognition of the altered states of the trimer observed here, we cannot conclude whether 10-1074 directly competes with Ab1456 and Ab1271, or rather competes allosterically (i.e., by trapping a closed conformation that is not accessible to Ab1456 or Ab1271). We note the human linear V3 mAb 3074^48^ exhibited a similar neutralization profile as that of Ab1456:^15^ e.g., both neutralized 6535.3, JRCSF, and X1632 strains to a greater degree than other strains.^48^ Given that mAb 3074 recognizes linear V3 peptides^44^ and preferentially binds Env trimers in the presence of soluble CD4,^35^ the finding of a vaccine-elicited monoclonal or polyclonal antibody response with a mAb 3074/Ab1456-like neutralization profile suggests targeting of undesirable open Env conformations, which is supported by our cryo-EM structures of Ab1456 in complex with open Env trimer conformations.

At least four [Ab1303,^12^ Ab1573,^12^ Ab1456 (this study), and Ab1271 (this study)] of the mAbs that we isolated after boosts 3 and 4 from NHPs immunized sequentially with SOSIP-VLP immunogens^15^ do not target conventional closed, prefusion Env trimer conformations, in contrast to the closed Env conformations in Fab-Env complexes involving mAbs isolated after the prime and first boost.^14,15^ Thus, it is interesting to consider how such antibodies were elicited. One possibility is that these antibodies were raised in response to SOSIP trimers with a propensity to sample open conformations. Consistent with this idea, heterologous serum neutralization titers from the animals that produced Ab1456 and Ab1271 spiked after receiving boost 2 with a B41-based SOSIP Env,^15^ a SOSIP that has been characterized as exhibiting more lability than BG505-based SOSIPs.^49,50^ Importantly, neutralization activity following the B41 boost increased against JRCSF, a strain that is neutralized by linear V3 and CD4i antibodies,^34^ suggesting that the B41-based boost 2 was responsible for eliciting Ab1456 / Ab1271-like responses. Alternatively, contaminating Env protomers may have elicited these antibodies, and only through the ability of these antibodies to also bind intact trimers, albeit in non-conventional closed conformations, can they neutralize a virus. In support of this idea, we previously showed by negative stain electron microscopy-based polyclonal epitope mapping (nsEMPEM)^51^ that antibodies elicited by our vaccine regimen caused SOSIP trimers to dissociate into protomers.^15^ Preventing SOSIP trimer immunogens from opening through the incorporation of an engineered disulfide (e.g., DS-SOSIPs) should limit exposure of epitopes exposed on open Env conformations^52,53^ but SOSIPs that have dissociated into protomers could also present Ab1456-like epitopes to the immune system. Therefore, preventing both Env trimer opening and trimer dissociation in vivo seem critical for avoiding Ab1456-like responses.

Antibody feedback mechanisms^54–58^ could also skew immune responses towards targeting non-closed conformations. While antibody feedback is typically thought of in terms of elicited antibodies forming immune complexes that shield targeted epitopes,^54–58^ some antibodies may also trap particular Env conformations; e.g., open Envs, and thereby facilitate further responses to non-closed Env conformations. Indeed, in each of the two NHPs from which these antibodies were isolated, we identified both a CD4bs mAb (Ab1573 or Ab1303) and a V3-targeting mAb (Ab1456 or Ab1271) that recognize non-closed Env conformations: Ab1573 and Ab1456 were isolated from NHP DGJI; Ab1303 and Ab1271 were isolated from NHP T15.^15^ Transitions of Env from closed to open conformations, Env trimer dissociation, and antibody feedback need not be mutually exclusive ways to elicit antibodies against non-closed Env conformations; thus, future studies to better understand mechanisms by which anti-Env antibodies are elicited are warranted.

The conformations of Env recognized by Ab1456 also raise questions regarding Env dynamics. For example, even in the absence of CD4, SOSIPs can sample open Env states and conformations other than that of a closed, prefusion trimer.^12,59^ SOSIP trimers, as well as Env trimers on virions, may transiently sample states such as the one observed in which only one of the three constituent protomers adopted a CD4-bound open conformation or an HT2-like state in which two of three protomers adopted a CD4-bound open conformation in the absence of a bound ligand (Figure 2A). When a mAb such as Ab1456 recognizes one of these states, the trimer likely cannot return to a closed conformation without the antibody first dissociating. With Ab1456 Fab bound to an open protomer, the trimer may then transiently sample more “fully-open” states in which all three protomers adopt CD4-bound open conformations (Figure 2A). Our multiple structural classes of the JRCSF SOSIP in complex with Ab1456 Fab(s) hint at different pathways for progressive trimer opening. Of note, we identified a single structural class of one Fab-bound JRCSF Env in which the trimer adopted an HT2-like state. In this conformation, protomer A (the bound protomer) as well as a neighboring unbound protomer (protomer B, clockwise to protomer A) both adopted a CD4-bound open conformation. Notably, we did not observe a structural class in which protomer B, but not protomer A, was bound by Ab1456 Fab, potentially suggesting that protomers on an HIV-1 Env trimer open in a clockwise fashion.

Previous studies showed that the duration of virus and antibody preincubation can influence the IC_50_ values of antibodies that neutralize HIV-1 by particular mechanisms; e.g., increasing preincubation times resulted in increasing potencies (decreasing IC_50_ values) of antibodies that neutralized by accelerating trimer decay.^60^ Here, we tested whether omitting virus and antibody preincubation would reduce the potency (i.e., increase IC_50_ values) of antibodies that recognize open Env conformations. This was not the case, however, suggesting that such antibodies do not have to wait for, and then capture, trimers as they sample various open conformations. Instead, it seems likely that viruses that are sensitive to Ab1456 and antibodies with similar recognition properties are already displaying their Envs in open conformations and that the neutralization profile of Ab1456, which mostly neutralizes tier 1 viruses with the exception of the tier 2 virus JRCSF,^15^ is an indicator of the conformational flexibility of these Envs.

The 6535.3 SOSIP Env trimer structures, both bound by Ab1271 and unliganded, raise further questions. For example, it will be important to determine whether the apex-altered conformation that we observed with the 6535.3 SOSIP is related to this Env being derived from a tier 1 HIV-1 and/or whether this state can also be seen in native, virion-embedded Envs. In any case, this unusual apex conformation may result from Env protomers that are transitioning to a CD4-bound conformation. Finally, it will be important to ascertain to what extent Envs from different HIV-1 strains exhibit differing dynamics at the trimer apex since this may influence their utility as potential immunogens.

Our structural analyses of Ab1456 and Ab1271 highlight the value of single-particle cryo-EM for deciphering underlying heterogeneities that can be present in antibody-antigen interactions. For example, Ab1456 Fab bound to the JRCSF SOSIP with varying stoichiometries, recognized different structural states of the trimer, bound to different configurations of protomers on a trimer, and even bound Env trimers with different angles of approach, all of which were resolved as different structural classes by cryo-EM. Although understanding this degree of heterogeneity is important for understanding the many ways in which Ab1456 can recognize Env, it limited the resolution of our 3D reconstructions, in part by reducing the number of particles in each class. Despite extensive classification, it is likely that heterogeneity persisted within the constituent particles of a given 3D class. By collecting larger datasets, it might be possible to determine higher-resolution structures and potentially even identify different structural classes. However, our reported structures were of sufficient resolution for observing V1V2 rearrangements and outward gp120 rotations, providing important insights into the Env conformations being targeted, as well as revealing epitope information.

Both Ab1456 and Ab1271 appear to recognize epitopes that are not available on closed Env trimers; therefore, both Env dynamics and recognition of specific epitope residues may influence antibody binding. For example, an Env trimer may include residues recognized by the antibody, but rarely, if ever, sample Env conformations that expose these epitopes for recognition. Conversely, an Env may sample the conformation recognized by an antibody (e.g., a CD4-bound open conformation for Ab1456), but lack sequence and or structure requirements for antibody recognition. The complex interplay between these factors likely limits the utility of antibodies whose binding is constrained by the conformational availability of the epitope; thus, consideration should be given to whether immunogens could be designed to avoid eliciting such antibodies.

In summary, here we examined the structure and function of two NHP mAbs that were elicited in a sequential SOSIP-based immunization approach, yielding results that rationalize the limited ability of heterologously neutralizing antibodies induced by this vaccine regimen to protect from a SHIV challenge.^15^ Although both mAbs exhibited heterologous neutralization breadth and targeted the V3 region of HIV-1 Env as intended,^14,15^ the mAbs also bound Env trimers in conformational states distinct from a typical closed, prefusion trimeric SOSIP conformation. Thus, an important finding of our present and previous analyses^14,15^ is that the appearance of heterologous neutralization breadth does not necessarily predict the presence of emerging bNAb lineages. Instead, we suggest that heterologous breadth elicited by a vaccine regimen should be examined in comparison to neutralization profiles of undesirable anti-Env antibodies to determine whether the stimulated antibody lineages represent a dead end or have the potential to mature along desired pathways. Of particular relevance to immunogen design efforts, the appearance of a neutralization profile consistent with CD4i or linear V3 antibodies, involving heterologous activity only against tier 1 HIV-1 strains and/or JRCSF, may indicate recognition of an undesirable epitope and consequently the inability to mature into a functional bNAb capable of robust protection from HIV-1 infection.

## Methods

### Protein Expression and purification

JRCSF (JRCSF.JB) and 6535.3 Envs were expressed as soluble chimeric SOSIP trimers^16^ (i.e., comprising a gp120 from JRCSF or 6535.3 paired with a BG505 gp41 and including stabilizing MD39 substitutions in gp41^30^). Relevant gp120 genes were synthesized (IDT gBlocks^TM^) and subcloned into a pcDNA3.1 expression plasmid backbone containing a gene encoding the stabilized BG505 gp41. SOSIPs were expressed via transient transfection of Expi293F cells with a 4:1 ratio of SOSIP- and soluble furin-encoding plasmids and then purified from transfected cell supernatants by immunoaffinity chromatography using immobilized mAbs (PGT145 for JRCSF or 2G12 for 6535.3) followed by SEC as described.^61^ Soluble Env trimers were stored at 4°C in 20 mM Tris pH 8.0, 150 mM NaCl (TBS).

Previously-reported IgG mAbs^15^ used in this study were expressed via transient transfection of Expi293 cells as chimeric IgGs with NHP V_H_-V_L_ domains and human IgG1 constant regions and purified by MabSelect SuRe chromatography (Cytiva). To produce Fabs for structural and biochemical experiments, IgG antibodies in phosphate-buffered saline (PBS) were cleaved by papain digestion using activated crystallized papain (Sigma-Aldrich) for 30 to 60 min at 37°C at a 1:100 enzyme:IgG ratio. Digested protein was applied to a 1 mL HiTrap MabSelect SuRe column (Cytiva) and flowthrough containing Fabs was collected. Fabs were further purified by SEC in TBS using a Superdex 200 Increase 10/300 column (GE Healthcare Life Sciences) before concentrating and storage at 4°C.

### Single-particle cryo-EM

For single-particle cryo-EM, SOSIP was complexed with Fab at room temperature, overnight with an approximate 1.3:1 molar excess Fab: SOSIP protomer. Samples were purified on a Superose 6 Increase 10/300 column (GE Healthcare Life Sciences) operating in TBS and leading SEC fractions were further concentrated to ∼1.1 mg/mL (JRCSF / Ab1456 sample) or ∼1.9 mg/mL (6535.3 / Ab1271 sample) using 10 kDa spin concentrators (Millipore). The samples were supplemented with octyl-maltoside, fluorinated solution (Anatrace) to a final concentration of 0.02% (JRCSF / Ab1456 sample) or 0.01% (6535.3 / Ab1271 sample) immediately before deposition of 3 µL onto a 300 Cu mesh, Quantifoil R1.2/1.3 grid (Electron Microscopy Sciences) that had been glow discharged for 1 min at 20 mA using a PELCO easiGlow (Ted Pella). Using a Mark IV Vitrobot (Thermo Fisher), the samples were blotted with a blot force of 0 for 3 s using Whatman No. 1 filter paper at 22C and 100% humidity and vitrified in liquid ethane.

### Data collection and Processing

40-frame movies were collected in super-resolution at a pixel size of 0.416 Å (105,000x magnification) using SerialEM^62^ on a 300 kV Titan Krios microscope (Thermo Fisher Scientific). Movies were collected using a 3x3 beam image shift with 3 shots per hole using beam-tilt compensation. The microscope was equipped with a K3 6k x 4k direct electron detector (Gatan) and a BioQuantum Energy Filter (Gatan) with a slit width of 10 eV. Collection parameters are described in Table S1 and the data processing workflow is shown in Figure S2 and S3. Briefly, movies were binned and patch motion corrected using CryoSPARC Live.^31^ The final particle stack used was picked using Topaz.^63^ Ab1271 / 6535.3 SOSIP data were initially processed by ab initio reconstruction and refinement in CryoSPARC.^31^ Particles were imported to RELION^33^ and subjected to 3D classification using the low-pass filtered, refined map as a reference model. Particles from select 3D classes were re-extracted in CryoSPARC^31^ and subject to ab initio reconstruction and non-uniform refinement^64^ to produce final maps. A similar workflow, but including iterative rounds of 3D classification, was used to process the Ab1456 / JRCSF SOSIP dataset.

### Mass Photometry

Mass photometry was performed on a OneMP (Refeyn). Glass coverslips (VWR) were pre-cleaned in water and isopropanol prior to use. Fab-SOSIP complexes were formed at room temperature in TBS overnight, at 4 µM trimer concentration and a 1.05 molar excess of Fab to SOSIP protomer. Fab-SOSIP complexes were diluted to 8 nM (trimer concentration) and diluted 10-fold (6535.3 SOSIP and 6535.3 SOSIP + Ab1271 Fab) or 4-fold (JRCSF SOSIP and JRCSF SOSIP + Ab1456 Fab) on the instrument. Movies were recorded for two minutes using Acquire^MP^ (Refeyn, v2023 R1.1) and analyzed using Discover^MP^ software (Refeyn, v2023 R1.2). A mass standard curve was prepared using beta amylase from sweet potato (dimer and tetramer). Figures were prepared on the Discover^MP^ software.

### Neutralization assays

The neutralization capacities of Ab1456 and other mAbs were assessed using TZM-bl reporter cells as described,^15^ including or not including the standard 1 h antibody-virus incubation step. Briefly, 96-well plates were seeded with TZM-bl cells (15,000 cells per well) overnight in Dulbecco’s modified Eagle’s medium (DMEM) containing 10% fetal bovine serum (FBS) and 100 U/mL Penicillin-Streptomycin-Glutamine (Gibco). For the standard assay, serial 5-fold dilutions of mAbs (e.g., 500, 100, 20, 4, 0.8, 0.16, 0.032, 0.0064, 0.00128 μg/mL) were incubated with transfection-derived or PBMC propagated virus at a multiplicity of infection (MOI) of 0.3 in a total volume of 100 μL in the presence of DEAE-dextran (40 μg/mL) for 1 hour at 37°C, and this mixture was then added to TZM-bl cells. For the modified test, mAb dilutions were first added to the cells in a volume of 50 μL, followed by the addition of virus in a volume of 50 μL, both in the presence of DEAE-dextran (40 μg/mL). TZM-bl cells were analyzed for luciferase expression 48 hours after virus addition using a Synergy Neo2 Multimode Microplate reader (Bio-Tek) with Gen5 version 1.11 software. Uninfected cells were used to correct for background luciferase activity. The infectivity of each virus without antibodies was set at 100%. The 50% inhibitory concentration (IC_50_) is the antibody concentration that reduces by 50% the relative light units (RLUs) compared with the no Ab control wells after correction for background. Nonlinear regression curves were determined and IC_50_ values calculated by using variable slope (four parameters) function in Prism software (v8.0).

Pseudoviral and SHIV stocks were generated by transfection of HEK293T cells. Briefly, 100-mm tissue culture dishes were seeded with 4x10^6^ HEK293T cells overnight in DMEM containing 10% FBS and 100 U/mL Penicillin-Streptomycin-Glutamine. Cells were transfected by adding 0.5 mL of a preincubated DMEM solution containing 4.5 μg of HIV-1 (SG3Δenv) backbone plasmids and 1.5 μg of HIV-1 Env plasmids or 6 μg of SHIV DNA, and 18 μL of FuGENE 6 transfection reagent (Promega), according to manufacturer’s recommendations. The cells were incubated at 37°C in a CO_2_ incubator for 48-72 h, and supernatant was harvested and stored at −80 °C in 0.5 mL aliquots. The generation of the rhesus PBMC-propagated SHIV_DH12-V3AD8_ challenge stock used for in vitro neutralization assays has been described.^43^

mAbs tested for neutralization were produced by co-transfecting paired heavy and light chain expression plasmids into Expi293F cells using ExpiFectamine 293 transfection reagents (ThermoFisher Scientific), purified from culture supernatants using the Protein A/Protein G GraviTrap kit (GE Healthcare), and buffer-exchanged into PBS as described.^65^

## Acknowledgements

We thank Songye Chen and the Caltech Cryo-EM Center, Anastasiya Oguienko, Morgan Abernathy, Welison Floriano, and Jost Vielmetter and the Caltech Protein Expression Center for experimental and technical support, and Malcolm Martin (NIH) for providing SHIV_DH12-V3AD8_ and SHIV_AD8-EO_ proviral DNA and the rhesus macaque PMBC-derived SHIV_DH12-V3AD8_ challenge stock.

This work was supported by the National Institute of Allergy and Infectious Diseases (NIAID) Grant HIVRAD P01 AI100148 (P.J.B., G.B.S., B.H.H.), R37 AI 150590 (B.H.H.), P01 AI 131251 (G.M.S.), NIH P50 1U54AI170856 (P.J.B.). This work was supported, in whole or in part, by the Bill & Melinda Gates Foundation grant INV-002143 (P.J.B.). Under the grant conditions of the Foundation, a Creative Commons Attribution 4.0 Generic License has already been assigned to the Author Accepted Manuscript version that might arise from this submission. A.T.D. was supported by an NSF Graduate Research Fellowship.

## Author contributions

Conceptualization, A.T.D., J.R.K., H.B.G., G.M.S., B.H.H., P.J.B.; Methodology, A.T.D., W.D., W.L.; Investigation, A.T.D., J.A.L., W.D., W.L., A.N.S.; Writing – Original Draft, A.T.D., B.H.H., P.J.B.; Writing – Review & Editing, A.T.D., J.R.K., H.B.G., J.A.L., A.N.S., B.H.H., P.J.B.; Visualization, A.T.D., W.D.; Supervision and Project Administration, J.R.K., G.M.S., B.H.H., P.J.B.; Funding Acquisition, G.M.S., B.H.H., P.J.B.

## Data and code availability

Cryo-EM maps of Ab1456 Fab / JRCSF SOSIP were deposited to the Electron Microscopy Data Bank (EMDB) and have accession codes: EMD-45944 (consensus structure), EMD-45945 (Class 1), EMD-45946 (Class 2), EMD-45947 (Class 3), EMD-45948 (Class 4), EMD-45949 (Class 5), EMD-45950 (Class 6), EMD-45951 (Class 7), EMD-45952 (Class 8). Ab1271 / 6535.3 SOSIP structures have accession codes: EMD-45942 (unbound) and EMD-45943 (bound). This paper does not report atomic models or original code. Additional information can be made available upon request.

## Supplemental Figures

**Figure S1.**
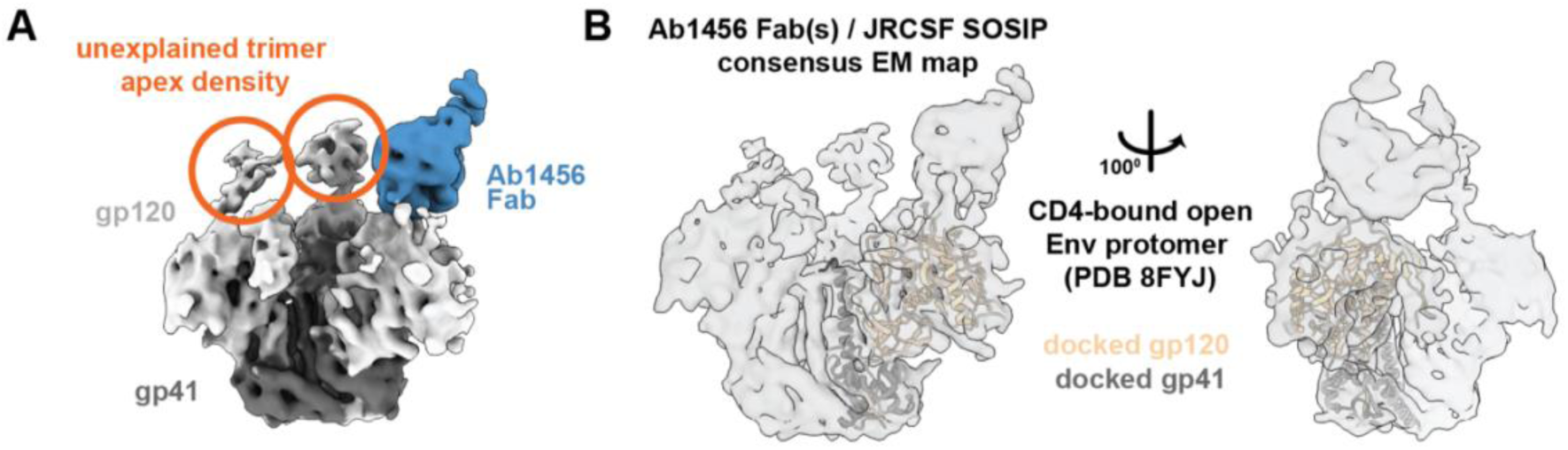
Analysis of Ab1456 Fab(s) / JRCSF SOSIP consensus map, related to Figure 2. (A) Unexplained density was present at the trimer apex of the consensus EM map. (B) Consensus EM density of the Ab1456 / JRCSF structure including a cartoon representation of docked coordinates of protomer A gp120 and gp41 from an Env heterotrimer (HT2) bound by two copies of soluble CD4 (PDB 8FYJ).

**Figure S2.**
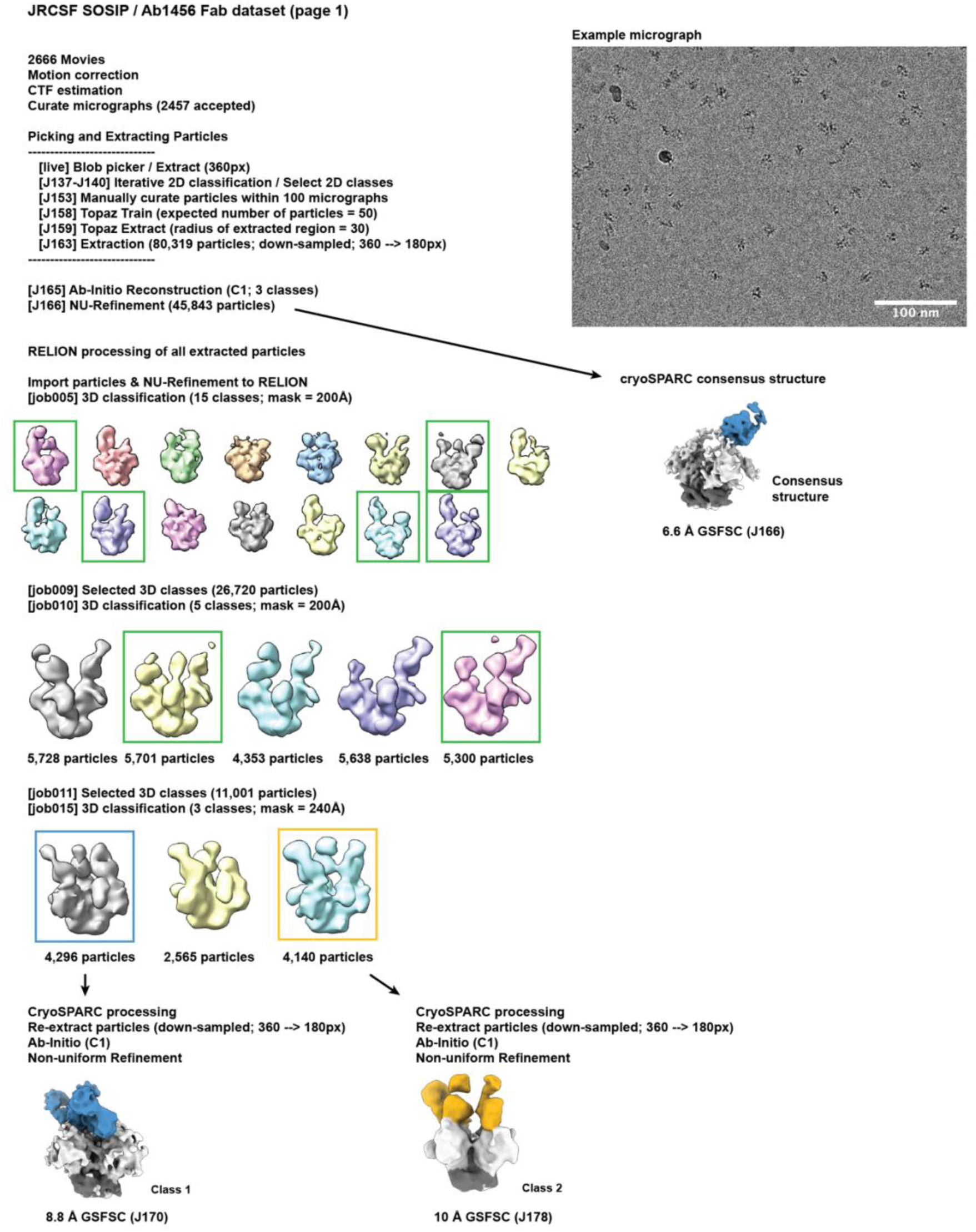
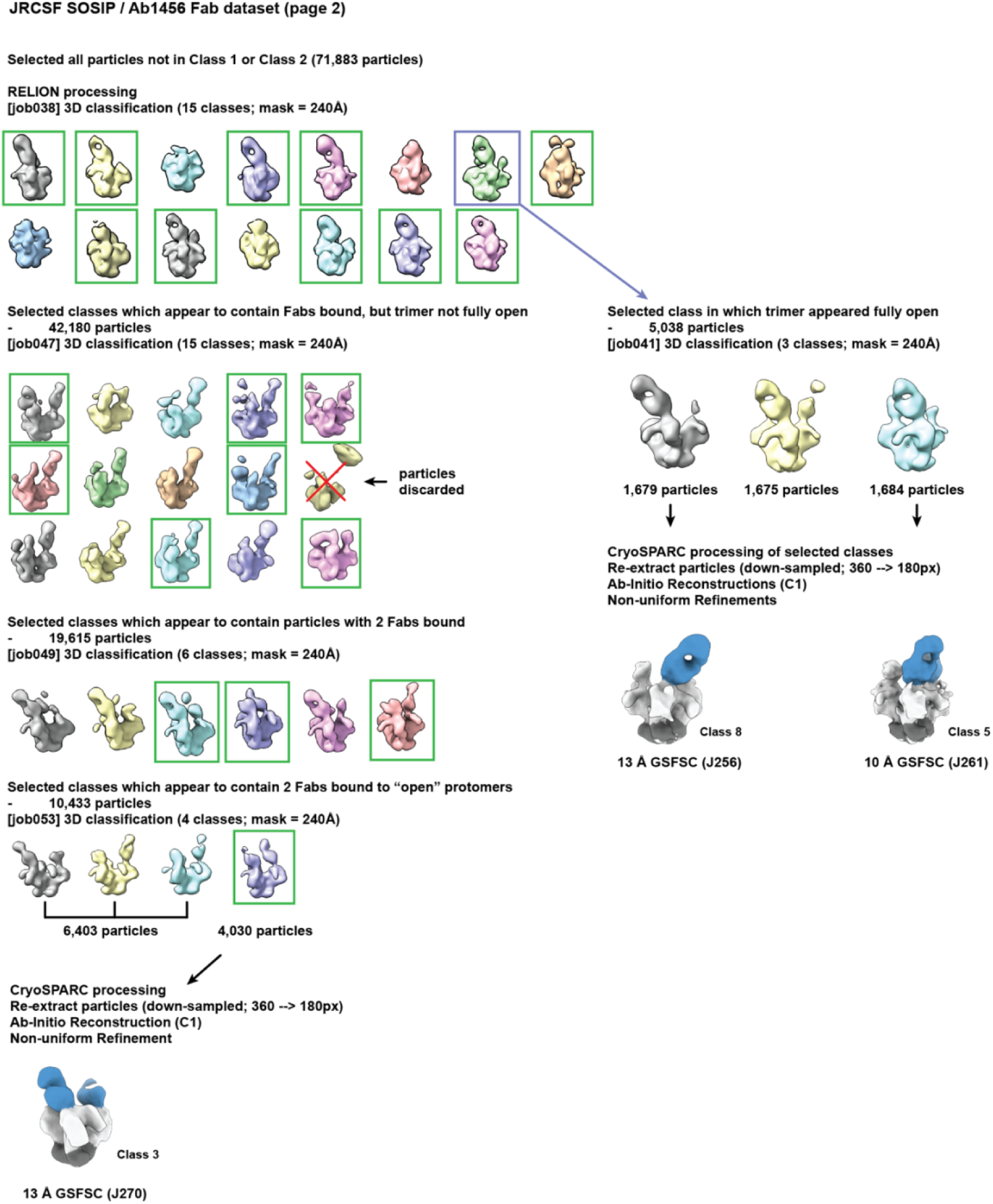
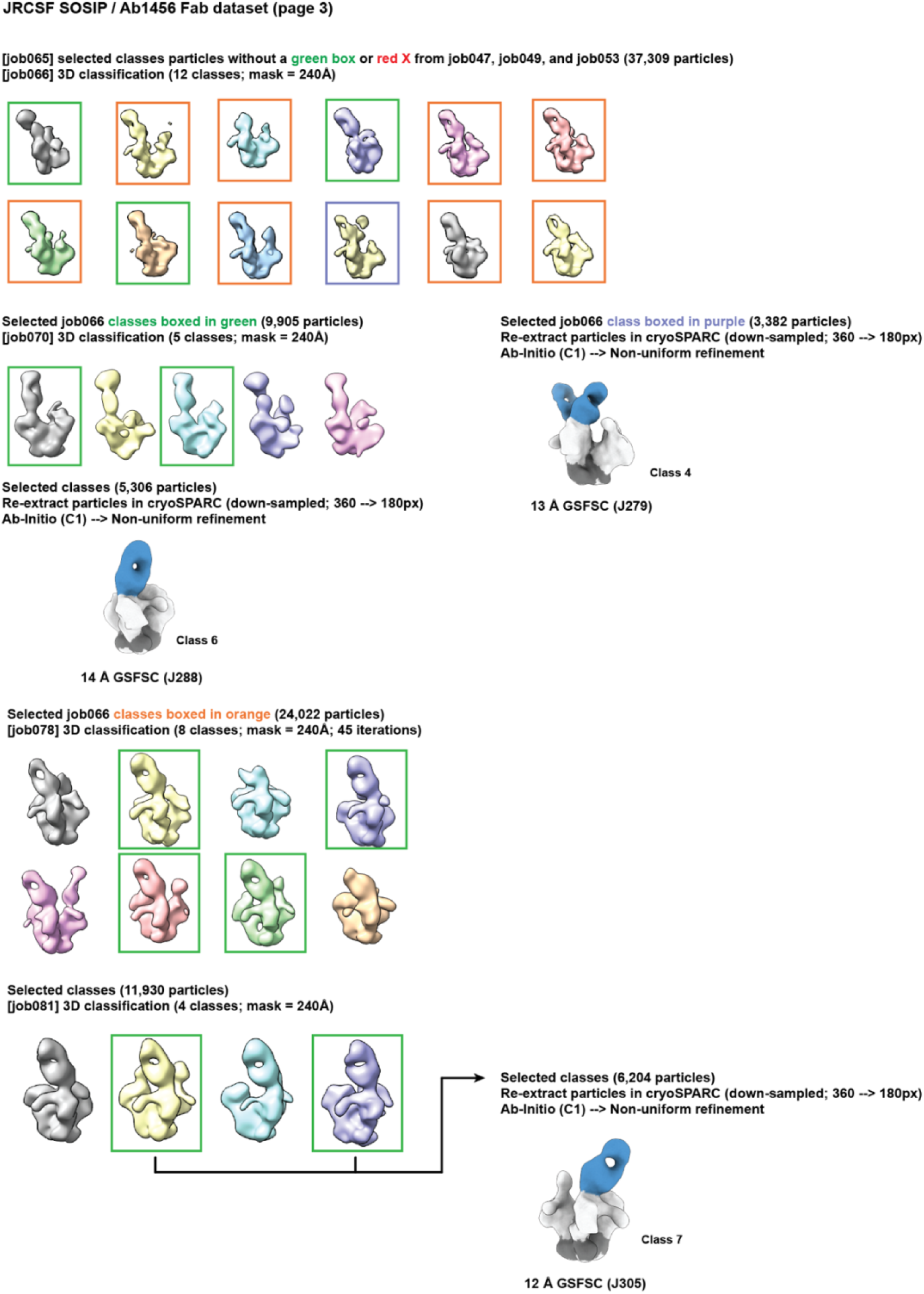
Data processing of the Ab1456 Fab / JRCSF SOSIP dataset, related to Figure 2.

**Figure S3.**
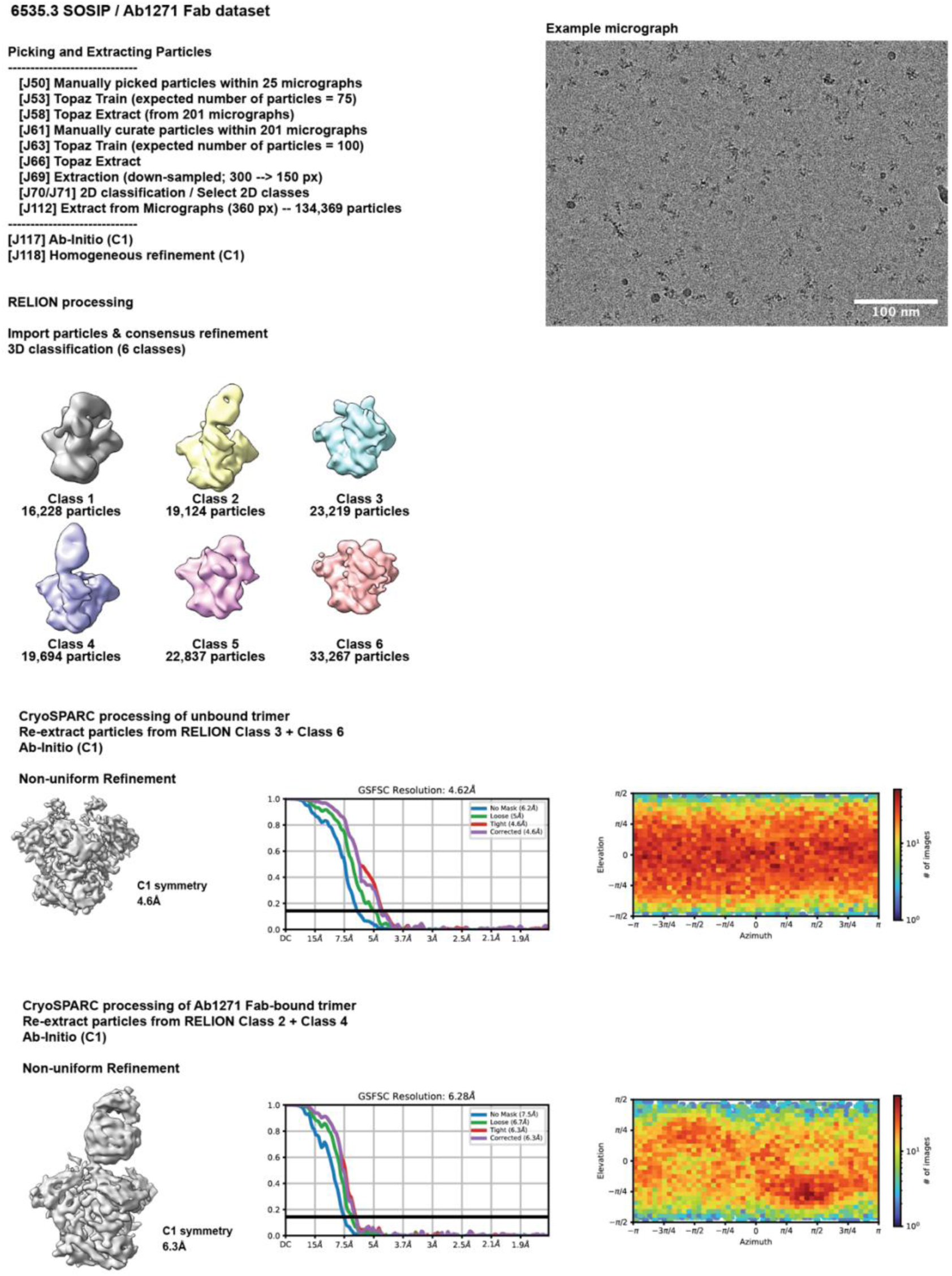
Data processing of the Ab1271 Fab / 6535.3 SOSIP dataset, related to Figure 3.

**Figure S4.**
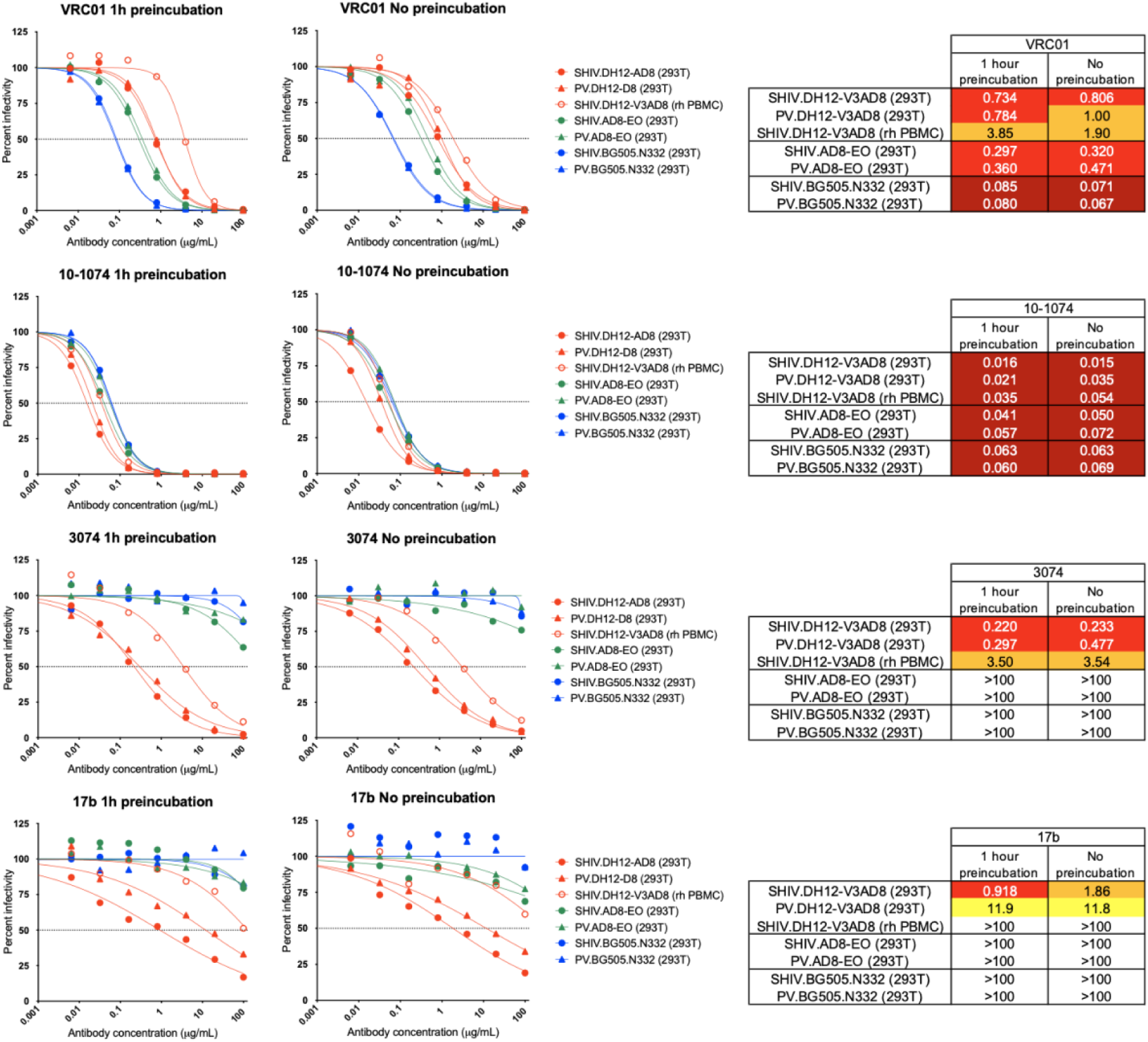
Omission of the antibody-virus preincubation step does not affect the potency of bNAbs or antibodies recognizing non-closed Env trimers, related to Figure 4. The sensitivity of viruses expressing the DH12-V3AD8 (red), AD8-EO (green) and BG505.N332 (blue) Envs to neutralization by VRC01, 10-1074, 3074, and 17b in a standard TZMbl assay including a 1 hour antibody and virus preincubation step^41,42^ and a modified assay with no preincubation are shown. Neutralization curves for the indicated mAbs are shown on the left (dashed lines indicate 50% reduction in virus infectivity), with the corresponding 50% inhibitory concentrations (IC_50_) in µg/mL shown on the right (coloring indicates relative neutralization potency). Pseudoviruses (PV) as well as replication-competent SHIVs derived either by HEK293T transfection (293T) or following propagation in rhesus macaque PBMC (rh PBMC) were tested. Note that similar to Ab1456 (Figure 4A), the slopes of the neutralization curves for the linear V3 mAb 3074 and the CD4-induced mAb 17b are more shallow slopes than slopes of the CD4bs bNAb VRC01^46^ and the V3 glycan bNAb 10-1074.^45^

**Table S1.**
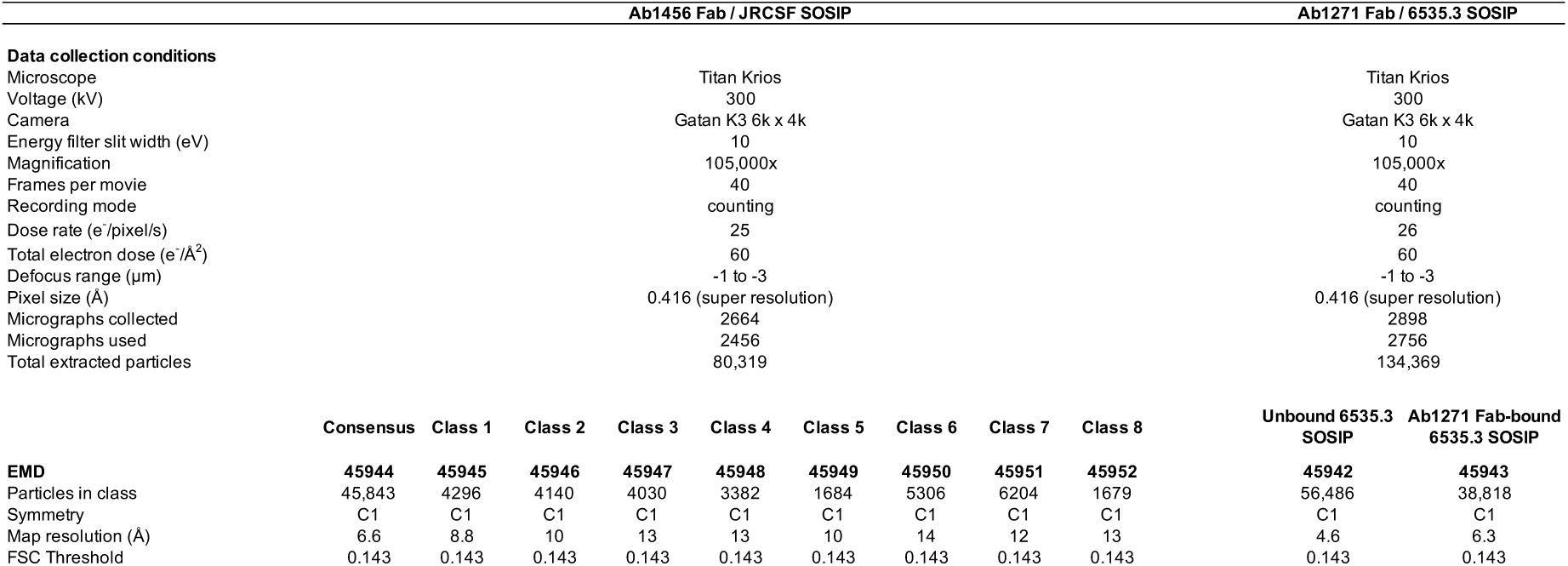
EM data collection and processing statistics.

